## Supplemental Data for "A role for the spinal cord cholinergic neuron circadian clock in RNA metabolism and mediating ALS disease phenotypes"

**SUPPLEMENTARY MATERIAL**

Supplemental Materials & Methods

Supplemental Figure S1 - Cre-Lox recombination enables genetic deletion of *Bmal1* in spinal cord cholinergic neurons.

Supplemental Figure S2 - GO enrichment analysis of oscillating genes in cholinergic neurons with peak expression during the light and dark periods.

Supplemental Figure S3 - Genes encoding ALS-linked RBPs are rhythmically expressed in mouse CNS regions.

Supplemental Figure S4 - Pathway enrichment analysis of differentially expressed genes between sALS patient LMNs and healthy controls.

Supplemental Figure S5 - Cholinergic neuron-specific *Bmal1*-deletion does not alter activity or RER rhythms, food intake, or weight.

**Uploaded as Supplemental Material online:**

Supplemental Tables S1–S8

### SUPPLEMENTAL MATERIALS & METHODS

*Activity, feeding, and weight measurements*

Mice were individually housed in Promethion Core metabolic cages (Sable Systems International) where they were allowed free access to food and water and acclimated for 4 days prior to data collection. Locomotor activity (measured by beam breaks), respiratory exchange ratio (RER) (measured by rate of carbon dioxide emission [$VCO_2$] divided by rate of oxygen consumption [$VO_2$]), food intake, and body weight were continuously monitored for 3 days. To obtain spinal cord measurements, fresh spinal cord tissue was isolated, weighed, and normalized to mouse whole-body weight.

*Lumbar spinal cord immunohistochemistry*

Mice were anesthetized with intraperitoneal injection of Euthasol (390mg/mL pentobarbital sodium, 50mg/mL phenytoin sodium), and then transcardially perfused with PBS followed by 4% PFA in 0.4M phosphate buffer (pH 7.4). Lumbar spinal cord was isolated, post-fixed in the same fixative overnight at 4°C, and paraffin embedded. 6µm thick sections were cut and mounted onto slides. Samples were then deparaffinized and hydrated prior to antigen retrieval by treatment with Antigen Decloaker (Biocare Medical, CB910M) in a high-pressure chamber for 20 min at 121°C (Deng et al. 2011). For immunohistochemistry, endogenous peroxidase activity was inhibited using 2% hydrogen peroxide. Samples were blocked for nonspecific binding by incubation with 1% BSA for 1 h at room temperature, and subsequently incubated with BMAL1 primary antibody (1:500 dilution; Novus Biologicals, NB100-2288) in a humidified chamber overnight at 4°C. The following day, samples were incubated with species-specific Alexa Fluor secondary antibody (1:500 dilution; Invitrogen) in the dark for 1 h at room temperature. Finally, samples were mounted with ProLong Gold Antifade Mountant (Invitrogen, P36930) and imaged on a DMI4000B confocal microscope (Leica Microsystems).

*Quantitative real-time PCR*

Following RNA isolation, cDNA was synthesized using the High-Capacity cDNA Reverse Transcription Kit (Applied Biosystems, 4368813). Quantitative real-time PCR (qPCR) was performed with iTaq Universal SYBR Green Supermix (Bio-Rad, 1725124) using a CFX384 Touch Real-Time PCR System (Bio-Rad) and analyzed using Bio-Rad CFX Manager Software (v3.1). Relative expression levels normalized to *β-actin* (steady between conditions) were determined using the comparative CT method. All primer sequences are listed in Table S8.

### Supplemental Figure S1

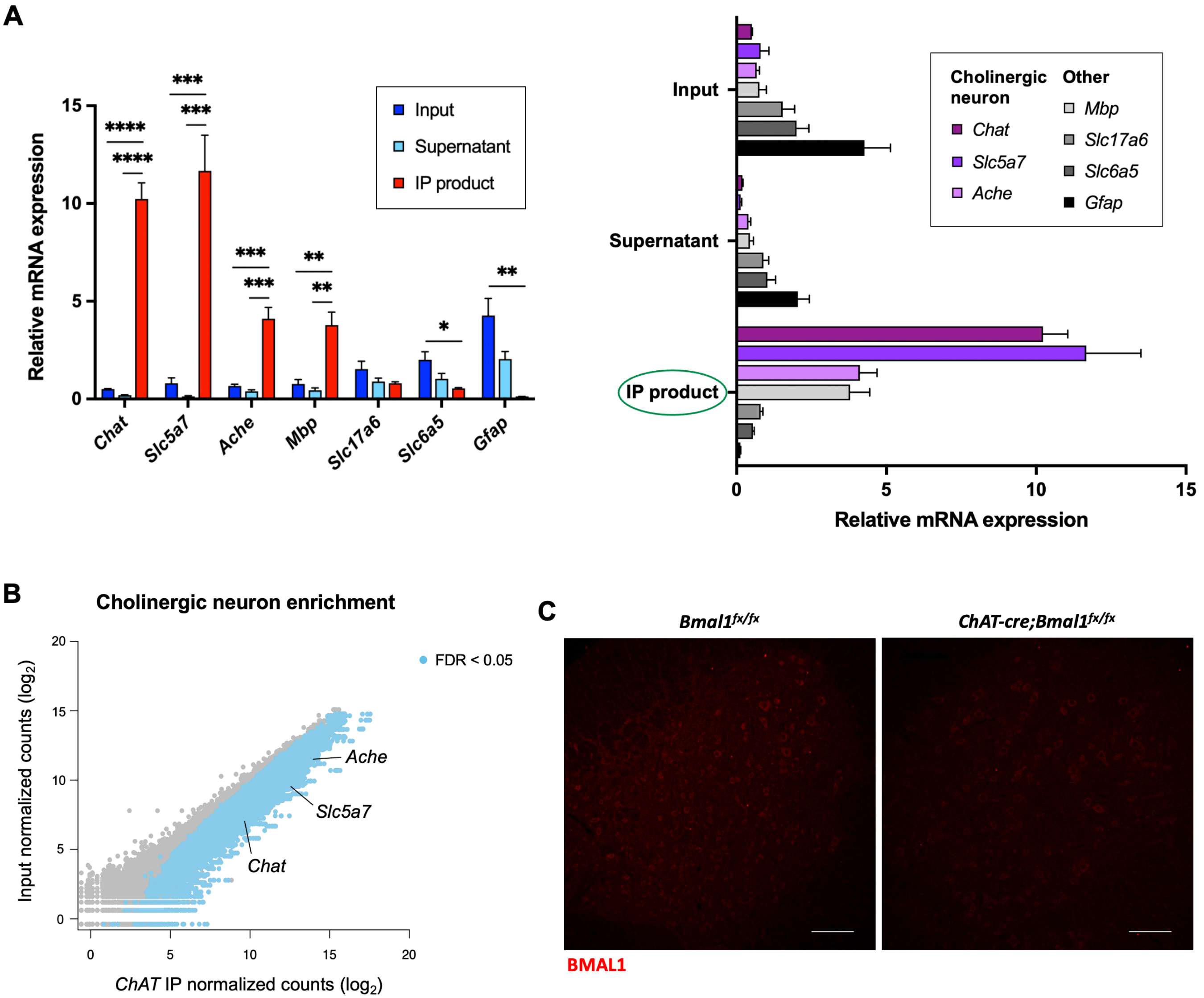


**Supplemental Figure S1. Cre-Lox recombination enables genetic deletion of *Bmal1* in spinal cord cholinergic neurons. (A and B)** Cholinergic neuron enrichment in WT *ChAT-Cre*;*RiboTag*$^{fx/+}$ male mice spinal cord IP product, as assessed independently by qPCR ($n$ = 3) (A) and DESeq2 normalized counts (B). Data are represented as mean ± SEM. Statistical significance was calculated by one-way ANOVA with Dunnett's correction for multiple comparisons ($^{*}p < 0.05$, $^{**}p < 0.01$, $^{***}p < 0.001$, $^{****}p < 0.0001$) (A) or the Wald test with Benjamini-Hochberg correction (B). **(C)** Representative immunofluorescence staining of BMAL1 (red) in lumbar spinal cord sections isolated from male mice with cholinergic neuron-specific *Bmal1*-deletion (*ChAT-Cre*;*Bmal1*$^{fx/fx}$ mice) and controls (*Bmal1*$^{fx/fx}$ mice). Scale bar represents 100µm.

### Supplemental Figure S2

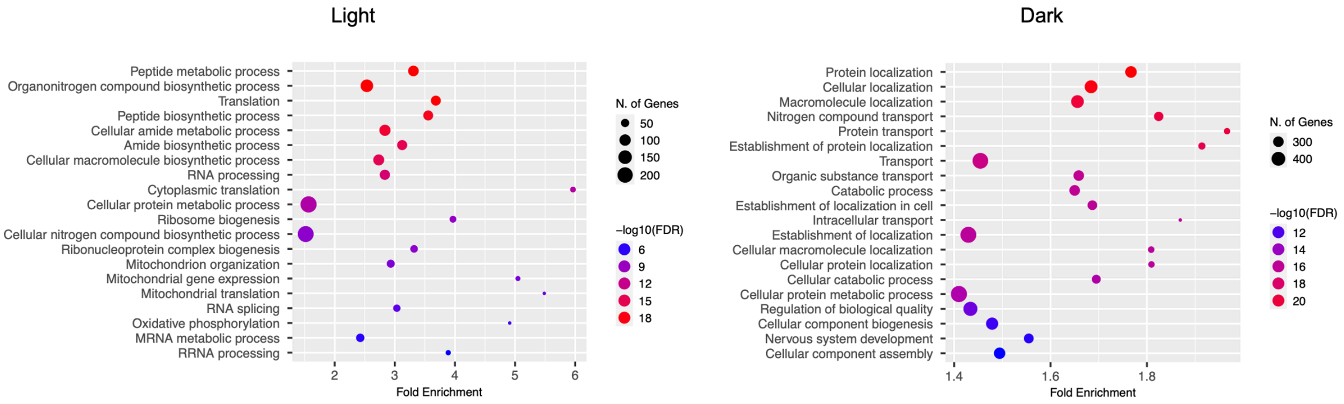


**Supplemental Figure S2. GO enrichment analysis of oscillating genes in cholinergic neurons with peak expression during the light and dark periods.**

**Supplemental Figure S3**

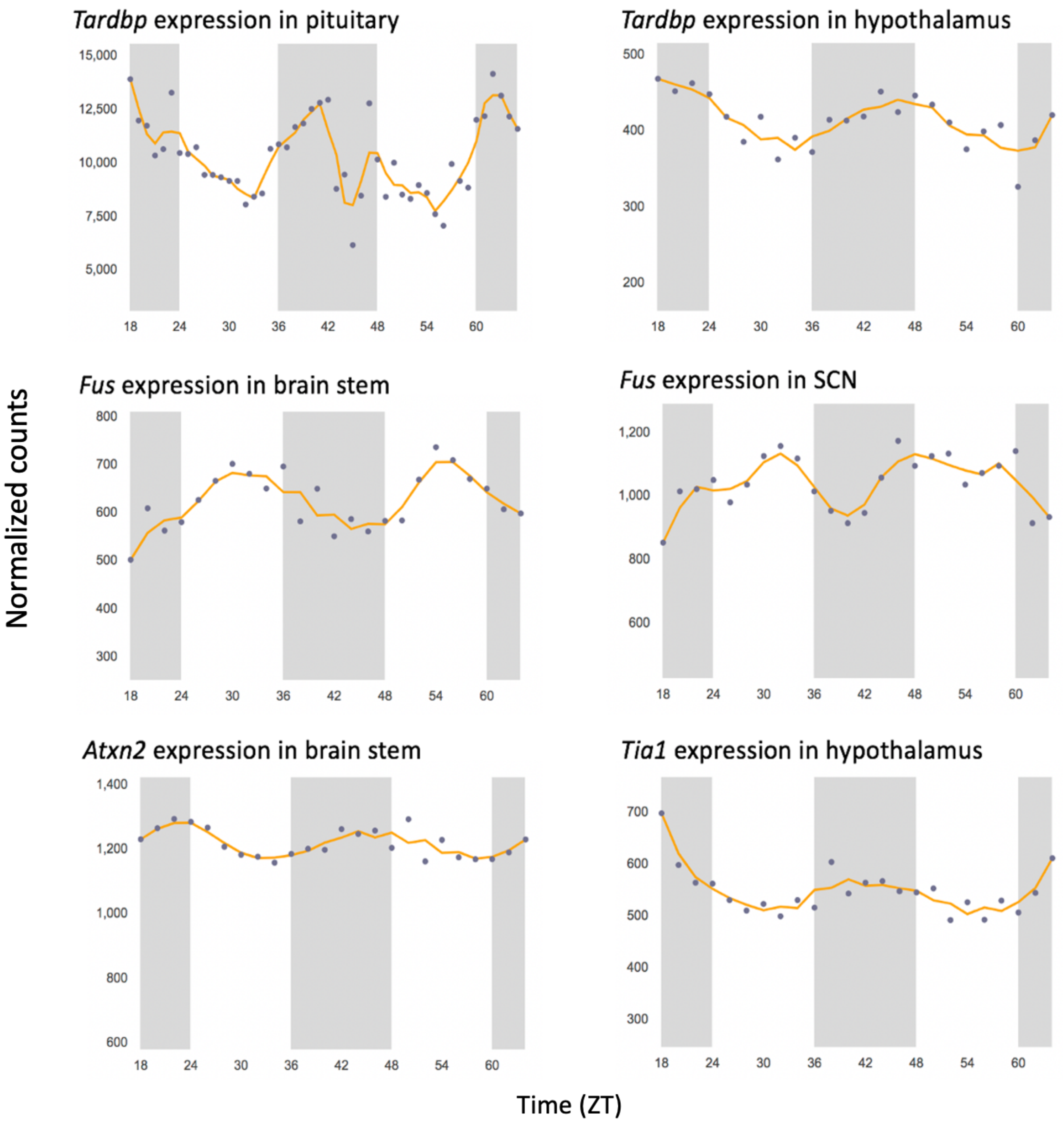


**Supplemental Figure S3. Genes encoding ALS-linked RBPs are rhythmically expressed in mouse CNS regions.** Rhythmicity was determined by JTK_Cycle analysis ($p < 0.01$). The shaded regions denote the dark/active time periods. Data obtained from CircaDB (Pizarro et al. 2013). CNS, central nervous system; SCN, suprachiasmatic nucleus.

**Supplemental Figure S4**

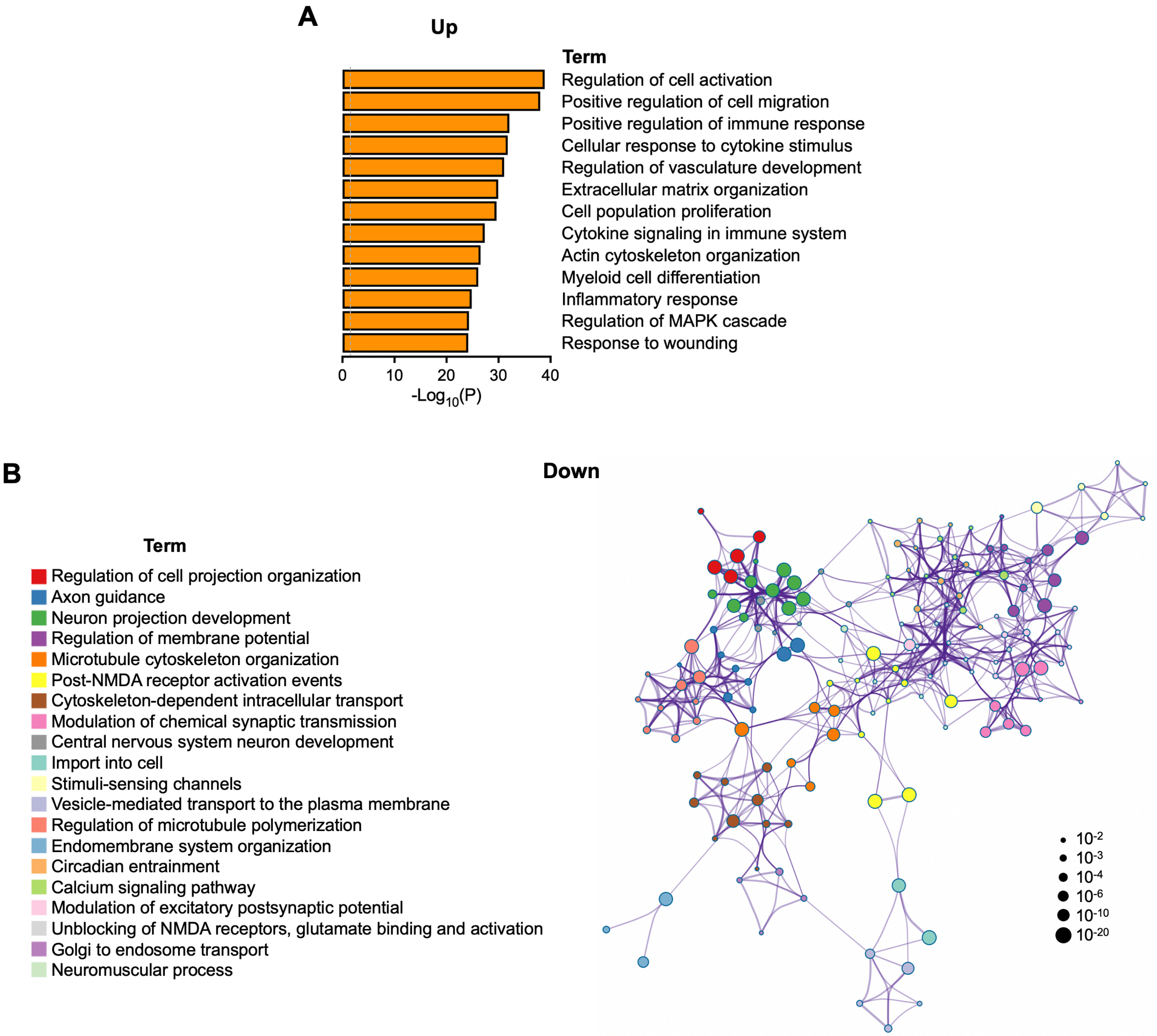


**Supplemental Figure S4. Pathway enrichment analysis of differentially expressed genes between sALS patient LMNs and healthy controls. (A)** KEGG, GO, and Reactome functional pathway analyses of genes that are upregulated in sporadic ALS (sALS) patient lumbar spinal cord motor neurons vs. healthy controls. **(B)** Network interconnection map showing functional pathway enrichment of genes that are downregulated in sALS patient lumbar spinal cord motor neurons vs. healthy controls. Circle nodes are color-coordinated by cluster identity. The size of each node is proportional to the number of genes within a given term. Terms with a similarity score > 0.3 are linked by an edge, with thicker edges reflecting higher similarity scores ($n$ = 13 sALS patients and 8 healthy controls). Re-analysis of data set from Krach et al. 2018. LMN, lower motor neuron.

Supplemental Figure S5

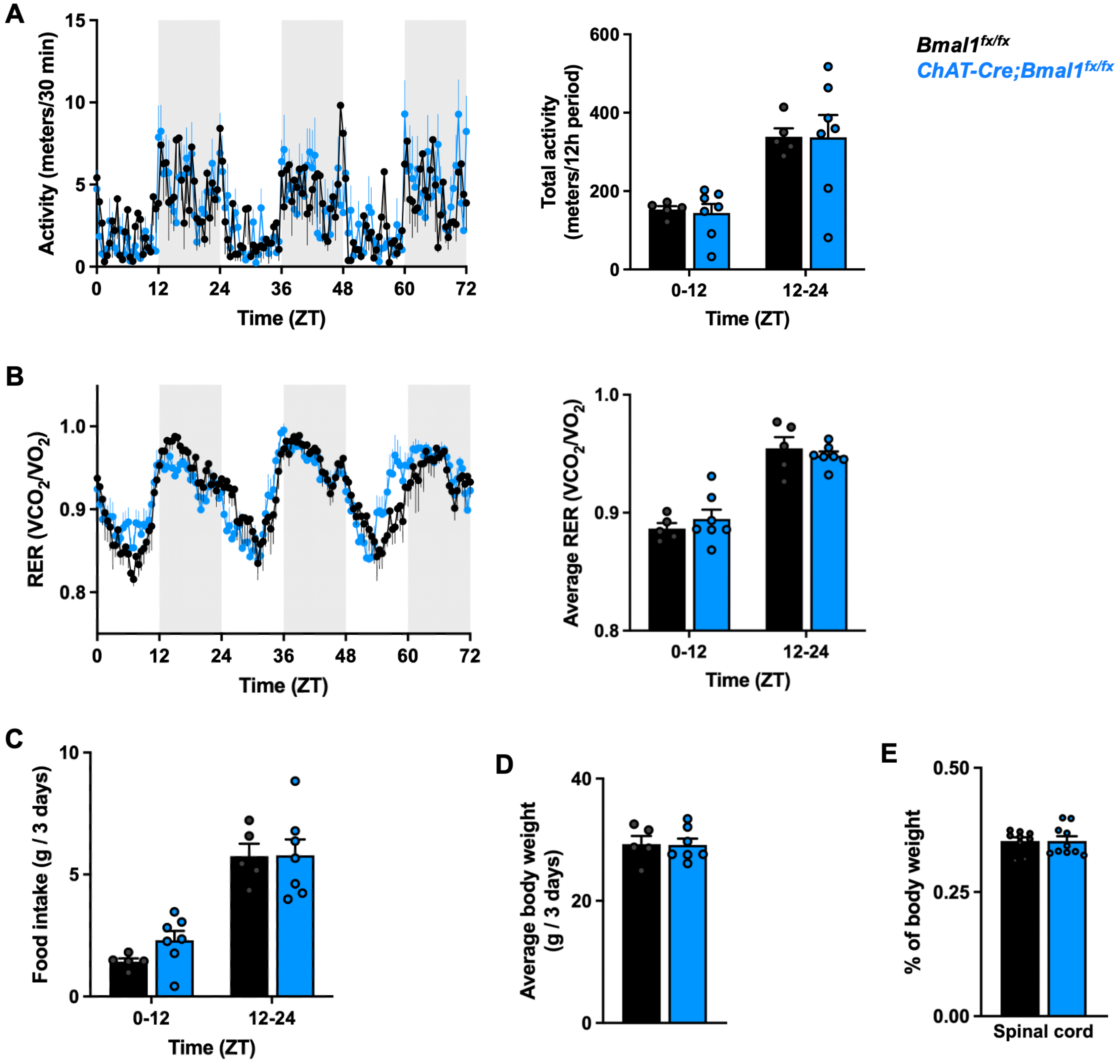


**Supplemental Figure S5. Cholinergic neuron-specific *Bmal1*-deletion does not alter activity or RER rhythms, food intake, or weight. (A and B)** Activity rhythms and total activity over 12 hours (A) and respiratory exchange ratio (RER) rhythms and average RER over 12 hours (B) in 5–7-month-old male mice with cholinergic neuron-specific *Bmal1*-deletion (*ChAT-Cre*;*Bmal1*$^{fx/fx}$ mice) and controls (*Bmal1*$^{fx/fx}$ mice) recorded over 3 days (*n* = 5-7). The shaded regions denote the dark/active time periods. **(C and D)** Cumulative food intake within each 12-hr period (C) and average body weight (D) over 3 days (*n* = 5-7). **(E)** Weight of isolated spinal cords as a percentage of total body weight (*n* = 9-10). Data are represented as mean ± SEM. Statistical significance was calculated by two-way ANOVA, with no differences between control and experimental groups within each 12-hr period and $p < 0.0001$ for all comparisons between light (ZT0-12) and dark (ZT12-24) periods (A-C), or unpaired two-tailed Student's *t*-test (D and E).
